# ProtParts, an automated web server for clustering and partitioning protein dataset

**DOI:** 10.1101/2024.07.12.603234

**Authors:** Yuchen Li, Carolina Barra

## Abstract

Data leakage originating from protein sequence similarity shared among train and test sets can result in model overfitting and overestimation of model performance and utility. However, leakage is often subtle and might be difficult to eliminate. Available clustering tools often do not provide completely independent partitions, and in addition it is difficult to assess the statistical significance of those differences. In this study, we developed a clustering and partitioning tool, ProtParts, utilizing the E-value of BLAST to compute pairwise similarities between each pair of proteins and using a graph algorithm to generate clusters of similar sequences. This exhaustive clustering ensures the most independent partitions, giving a metric of statistical significance and, thereby enhancing the model generalization. A series of comparative analyses indicated that ProtParts clusters have higher silhouette coefficient and adjusted mutual information than other algorithms using k-mers or sequence percentage identity. Re-training three distinct predictive models revealed how sub-optimal data clustering and partitioning leads to overfitting and inflated performance during cross-validation. In contrast, training on ProtParts partitions demonstrated a more robust and improved model performance on predicting independent data. Based on these results, we deployed the user-friendly web server ProtParts (https://services.healthtech.dtu.dk/services/ProtParts-1.0) for protein partitioning prior to machine learning applications.

**GRAPHICAL ABSTRACT:** 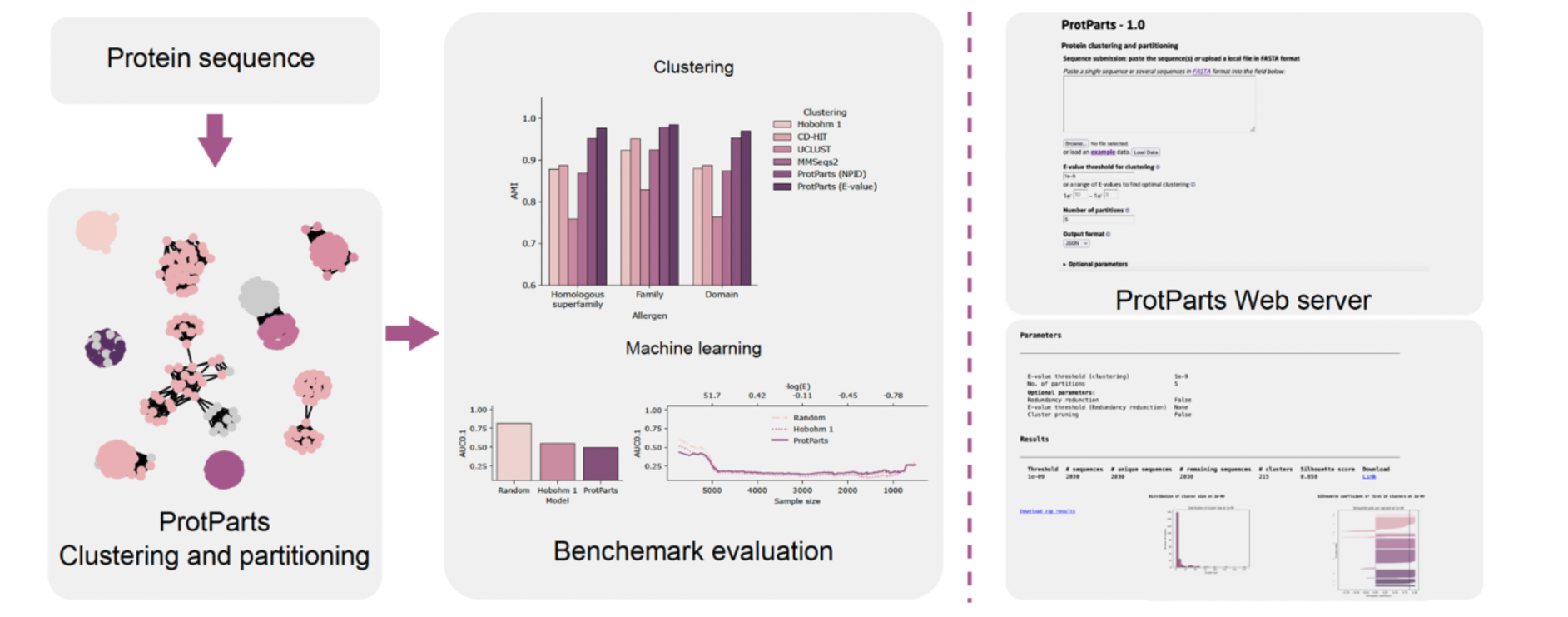

## INTRODUCTION

Machine learning methods have over the last decades been widely applied in bioinformatics to process and analyze protein sequences (1, 2). The capacity of accurately predicting protein structure and function using machine learning has largely accelerated the identification of sequence patterns and enrichment of the protein universe (3). In order to construct an effective machine learning predictive model, overfitting is an essential issue that cannot be overlooked. Here, overfitting refers to the situation where a model performs well on its training data but poorly on unseen evaluation data (4, 5). This lack of generalization ability can severely limit the future application of the developed models. One of the important factors that gives rise to overfitting is data leakage between the train and the test data, which is usually used to find the optimal hyperparameters of the model. The leakage of test data with similarities to the training datasets makes the models likely to memorize identical features instead of learning the underlying patterns. In bioinformatics, data leakage is pervasive and different solutions have been proposed to counteract it, for instance, clustered data splits on train and test have been used in protein properties prediction (6), protein-protein interaction prediction (7) and genomics (8). These studies have proved that the data leakage between training and test leads to a systematic overestimation of model performance and poor generalization. Due to their biological importance or research interest, certain proteins or protein families are extensively studied, and similar sequences constitute a large amount of sequences in protein databases (9, 10). Therefore, applying random partitioning on these imbalanced-distributed datasets to split training and test datasets is likely to create rhave edundancy between them, and leading to overfitting models with poor generalization abilities. Thus, addressing the sequence redundancy properly is imperative to avoid these issues.

Clustering categorizes data based on their similarity, significantly reducing data redundancy across partitions by dividing the clustered data into partitions. Early clustering tools have been developed to tackle the challenges for massive protein sequence clustering. CD-HIT, a greedy algorithm based on Hobohm (11), is the most commonly used tool in sequence clustering. It sorts sequences with a descending order and sets the longest sequence as the first representative sequence of the first cluster (12, 13). The following sequence on the sorted list is further compared to the representative sequence existing on the first cluster. If the query sequence falls below a defined threshold of percentage identity, it becomes a new representative sequence in a new cluster. Otherwise, it joins the cluster where it shares sequence similarity to its representative sequence. Due to the massive computation needed for sequence alignment, CD-HIT introduced k-mer filters to accelerate the clustering by skipping part of the full sequence alignment. The number of shared k-mers reflects the similarity between two sequences. Likewise, UCLUST applies a similar greedy algorithm but with a different alignment approach called USEARCH (14). USEARCH calculates the number of unique shared k-mers between query and target sequences, sorts sequences by the number of unique k-mers with a descending order and terminates comparison when the number of hits exceeds a preset threshold. MMseqs2 employs the greedy set cover algorithm, which iteratively generates sub-clusters by sequence percentage identity, E-value and coverage to find a representative sequence for each individual cluster (15, 16). Subsequently, it compares the member sequences with the representative sequences, merges the member sequences if they are similar above a threshold, and finally obtains the minimum representative sequences that cover the whole dataset. These clustering tools have been developed with the purpose of clustering and annotating large DNA datasets potentially including several genomes. However, the greedy algorithm does not necessarily find the optimal solution and may overlook similarities between member sequences. One sequence might not share similarity with representative sequences defining the clusters, yet it could still share similarity with other non-representative sequences inside a cluster. The unchecked sequence similarity has potential to introduce data leakage across clusters and partitions. This could impact the estimation of model performance in machine learning.

Additionally, these tools utilize k-mers to accelerate the sequence alignment and use them to define the sequence percentage identity for sequence clustering. The percentage identity only counts identically aligned amino acids, without taking the positively aligned amino acids into consideration. Amino acid substitutions are not random events; instead, amino acids are more frequently replaced by others that share similar chemical properties (17). In addition, different approaches to calculate the percentage identity of an alignment can have largely variant results (18, 19). Therefore, naively using percentage identity to compare a pair of sequences might lack biological relevance and statistical significance (20).

Trying to address the question of an exhaustive clustering specifically designed for protein datasets, we developed a user-friendly web server tool called ProtParts. Here, we compared sequence similarity measurements and investigated the clustering performance of different measurements and methods, proposing a new clustering method that provides better clustering, and generates partitions to avoid data leakage in using protein sequences. ProtParts clusters proteins in graphs and introduces E-value as the similarity calculations.

## MATERIAL AND METHODS

### Datasets

In Dataset 1, allergen proteins were collected from AllergenOnline v21 (September 2020), a curated database encompassing 2,171 IgE-binding validated antigens (either by IgE binding only -western blot, ELISA-without biological activity; or IgE binding plus biological activity test as measured by basophil activation or IgE plus Skin Prick test) (21). Proteins with inaccessible protein accession IDs, those containing non-canonical amino acid letters (B, J, Z and X), those having extreme sequence lengths (L < 50 or L > 1000), or duplicated sequences were excluded, resulting in a dataset consisting of 2,030 unique proteins. Additionally, protein labels, such as protein domain, family, homologous superfamily, were collected when available for all the allergens using the InterPro database (22). This smaller dataset consisting of 1001 sequences is referred as dataset 2. Furthermore, we collected the evolutionary relation of allergen species from the NCBI taxonomy common tree to investigate the allergen distribution across species (23). Additionally, 7,298 non-allergens were included using the same approach as in NetAllergen, where proteins from the same species were collected to reproduce the amino acid background frequencies of the allergens (24).

An independent allergen dataset (dataset 3) was collected to evaluate the classification performance of the different models produced in this work. This ‘evaluation dataset’ was retrieved from the evaluation dataset of NetAllergen (24), which originated from different allergen databases except AllergenOnline. The original evaluation dataset was filtered by removing identical sequences to those present in the allergen training dataset. This dataset contained 1,017 allergens and 4,715 non-allergens. The non-allergens in this dataset are proteins with no allergen description, from the same species as the allergens, and follow the same length distribution as the allergens.

Additionally, the ASTRAL SCOPe 2.08 (September 2021) dataset was used. This dataset is a manually curated dataset with domain, family and superfamily annotations from proteins of known structure in a hierarchy according to structural and evolutionary relationships (25, 26). This dataset contains 35,494 protein sequences that share a percentage identity of 95% or lower to other members of the dataset.

### Metrics for sequence similarity

Both the Allergen and the ASTRAL SCOPe datasets were individually searched in an all-against-all fashion using BLAST+ (2.12.0) with default parameters to obtain pairwise similarity metrics for clustering (27). Both percentage of identity and E-value were collected for comparing purposes. The E-value is a measure of the expected number of random hits given an alignment score and is proportional to the length of the sequences and the number of proteins in the database, and inversely and exponentially correlated to the alignment score. Therefore, a higher alignment score in the same database will provide a smaller E-value. For easier visualization purposes, in the plots, the E-value was converted to the negative logarithm of the E-value (-logE) with a base of 10. In this converted metric, a higher -logE value indicates a more similar sequence. Furthermore, the percentage identity provided by BLAST corresponds to the number of identical amino acids divided by the alignment length. Since the alignment length is a variable fraction in any pairwise alignment despite the length of the compared sequences, a ‘normalized percentage identity’ (NPID) was created, dividing the number of identical amino acids in the alignment by the length of the shorter sequence in the pairwise alignment. An identity of 50 matches on an alignment of 100 amino acids will therefore be different if the shortest protein sequence in the alignment is 100 amino acids long or 500 (28).

### Clustering methods

*ProtParts.* To cluster the sequences in each dataset, we created a graph in which every protein sequence is in a vertex and either the E-value or the percentage identity is used as the connecting edges representing protein similarity. The initial complete graph was constructed by connecting all proteins with similarity-weighted edges. Based on the weighted edges, we pruned connections exceeding an E-value similarity threshold, consequently the graph was split into multiple connected subgraphs and singleton subgraphs. The generated subgraphs became independent clusters and exhibited a similarity to the other clusters less than the given threshold.

*Hobohm 1*. We first sorted the dataset by decreasing sequence length, and in a similar fashion as the greedy incremental algorithm will do, we applied the Hobohm 1 algorithm, which takes the first sequence as the representative sequence in the first cluster (11). It next compares the second longest sequence to the first representative sequence. It could either be a member of the first cluster when the similarity metric is above a threshold, or a new representative sequence in the second cluster when it is below the defined similarity threshold. Each sequence on the ranked list is compared to each one the representative sequences of each cluster ordered by sequence length until it matches the similarity threshold, or it becomes the representative sequence of a new cluster. This process is iteratively repeated until the last and shortest sequence is clustered.

*Random*. Based on the clustering results of ProtParts and Hobohm 1, we set the number of clusters in random clustering approximately equal to the number of clusters in both ProtParts and Hobohm 1. Firstly, the number of clusters was determined, and empty clusters were initialized. Then, protein sequences were randomly assigned to initialized clusters.

### Classification tools

Three different prediction tools were applied in the training of the clustered allergen dataset. NetAllergen is a random forest model for allergenicity prediction with 60 features including immunological and structural information (27). DeepAlgPro encodes sequences as one-hot vectors and trains them using a convolutional neural network using an attention layer (29). ProPythia is a general tool for protein classification with machine learning and deep learning (30). In this study, ProPythia automated package was used to generate 1,997 descriptors (including amino acid composition, pseudo amino acid composition, physicochemical properties and autocorrelation) for sequences, and it was trained with a common feedforward neural network.

### Clustering evaluation metrics

The clustering performance was evaluated using two metrics: adjusted mutual information (AMI), and silhouette coefficient. Mutual information measured the amount of information contained in predicted cluster labels about the true labels. It can be regarded as the reduced entropy of identifying true labels having the knowledge of the predicted cluster label. High mutual information indicates more information about true labels is contained in predicted cluster labels. Here the true labels refer to the annotations of domain, family, and superfamily obtained from InterPro database on the allergen dataset 2, and to class, fold, family, and superfamily in the ASTRAL SCOPe dataset. To obtain the AMI, the mutual information was further adjusted to yield a score of one if the cluster labels fully overlap with the true labels, and a value of zero when cluster labels originate from random clustering (31). The adjustment eliminates the effect of having large number of clusters, which leads to high mutual information even though no additional information is shared.

Additionally, silhouette coefficient served as a means to compare different clustering algorithms. To compute the silhouette coefficient of a protein, it firstly calculated the average of similarity difference (cohesion) between the protein and all other proteins within the same cluster. Then it used the same approach to determine the average of similarity distance (separation) to all proteins in the nearest neighboring clusters. Finally, the silhouette coefficient was defined by subtracting the ratio of distance (cohesion) and distance (separation) from one. The overall average silhouette coefficient measured the consistency of protein similarities within the clusters.

We evaluated the performance of the models on the allergen dataset using the area under ROC curve (AUC) and AUC 0.1, which are metrics for binary classification. AUC is independent of the threshold selection and is not affected by the ratio of positive to negative samples.

AUC 0.1, a variation of the standard AUC, calculates the area under the curve up to a false positive rate of 0.1. AUC 0.1 specifically assesses how correctly models predict true positives while the false positive rate is low.

## RESULTS

### Dataset selection

To demonstrate the use of the clustering tools, we firstly collected 2030 allergen proteins from the AllergenOnline database (Dataset 1). These allergens exhibit both sequence diversity and redundancy, originating from 379 unique organisms. However, the allergens are not evenly distributed among the different species. For instance, the most frequent species, *Triticum aestivum*, comprises 4% of the allergens in Dataset 1, and most of these allergens share sequence similarity to each other. On the other hand, there are 158 species with only one allergen and each one merely account for 0.05% in the dataset. The high sequence similarity shared by many common allergens may come from the widespread use of sequence similarity to predict allergenicity, potentially biasing experimental assays used to determine it. These features allow the allergen dataset to simulate a common scenario in protein databases where numerous highly identical and unique sequences exist. Therefore, the application of ProtParts can be extended to more general cases of protein sequence clustering when high similarity is an issue.

For the analysis of the dataset and evaluation of the clustering results, structural and evolutionary related information of this dataset was collected from UniProt and InterPro. We expected that proteins sharing the labels: domain, family, or superfamily should belong to the same cluster, therefore we have used these labels as the ground truth to benchmark the clustering performance. Excluding proteins with none or only partial annotations for domain, family, and homologous superfamily, resulted in 1001 remaining allergens (Dataset 2) The Dataset 2 contained 78 unique or combined domains. With combined domains we refer here to the groups of proteins that have a combination of two or more domains. The most common domain on the allergen dataset was the bifunctional inhibitor/plant lipid transfer protein/seed storage helical domain (IPR016140), contributing with 19% allergens in Dataset 2. Allergens from *Triticum aestivum* species provided most of the data annotated to IPR016140. Additionally, the Bet v 1/Major latex domain (IPR000916), also known as part of the major birch pollen allergen, is present in around 18% of the allergens in Dataset 2. Interestingly, allergens from largely different species shared some of the domains (Supplementary Figure S1). Although they are quite far apart from a taxonomy perspective, this indicates these proteins are potential homologs and share remote relations. Therefore, this dataset constitutes a good example to investigate the effects of sequence similarity in machine learning applications and is suitable to assess proper clustering to avoid data leakage.

### Sequence similarity metrics

To determine sequence similarity in dataset 1, we used BLAST to query each protein against all the proteins in the dataset and obtained similarity metrics such as the percentage identity (PID), E-value, and the normalized percentage identity (NPID). Typically, clustering methods use PID or NPID to define thresholds for clustering. Here, we compared the NPID to the E-value, as the latter adds the information of what is the likelihood of the alignment happening by chance. The distribution of -logE-values and NPID demonstrated that they were positively correlated under different query lengths (Figure 1). It also suggested that longer sequences had a greater slope of E-values with respect to NPID. The alignment of shorter query sequences was mainly distributed at the lower right region where the -logE-values were approximately from -1 to 50 and the NPID was from 0 to 1. On the other hand, longer query sequences had the same range of NPID but a broader -logE-value from -1 to 180. Since the - logE-value increased more rapidly for longer query sequences, there were many alignments with a lower NPID but a significant E-value. Therefore using a typical identity threshold of 30% to define homologous proteins (20, 32, 33), the NPID or E-value did not always agree (Figure 1). To showcase the instances where the E-value and PID do not agree, we picked two alignments that were representative to the extreme situations. One where the E-value suggested homology but the percentage identity did not, and another case showing the opposite. For instance, the alignment between the allergen proteins A0555 (B3STU4, Car i 2) and A1046 (Q948Y0, Gly m 5) had an E-value of 3x10^-93^, a NPID of 24%, and query coverage of 52%. However, both of them are vicilin-like/vicilin proteins and also share the same Cupin 1 domain. Even though a strict cutoff of NPID of 30% suggested that these proteins would be unrelated, the very low E-value in the alignment indicated the two sequences were not aligned by chance, suggesting significant homology. To further investigate what led to the differences, we examined the BLAST alignment of these two sequences. In addition to the identical amino acids, the alignment had 265 (63%) positively aligned residues (aligned amino acids that may share functional properties) (Supplementary Figure S2A). On the contrary, there were few cases where the NPID over 30% suggested homology while the E-value was not significant. For example, the alignment of A1986 (P04724, alpha/beta gliadin) and A2044 (P04730, gamma gliadin) had an E-value of 4.6, NPID of 39% and query coverage of 81%, and they shared the seed storage helical domain. In this case, the NPID suggested a relationship of the two proteins, but the E-value did not. From the alignment of A2044 and A1986, we observed a short alignment, with a large number of repeated glutamines, and simpler amino acid compositions (Supplementary Figure S2B). In this case, the low complexity of the sequences suggested that it is possible to align sequences that are not necessarily related by chance. Importantly, this information could be captured by the non-significant E-value obtained for this alignment. Thus, based on these representative alignments, we can suggest that the E-value was more suited to reveal protein homology with high sequence complexity, while the NPID was able to find the low complexity sequence pairs.

**Figure 1.**
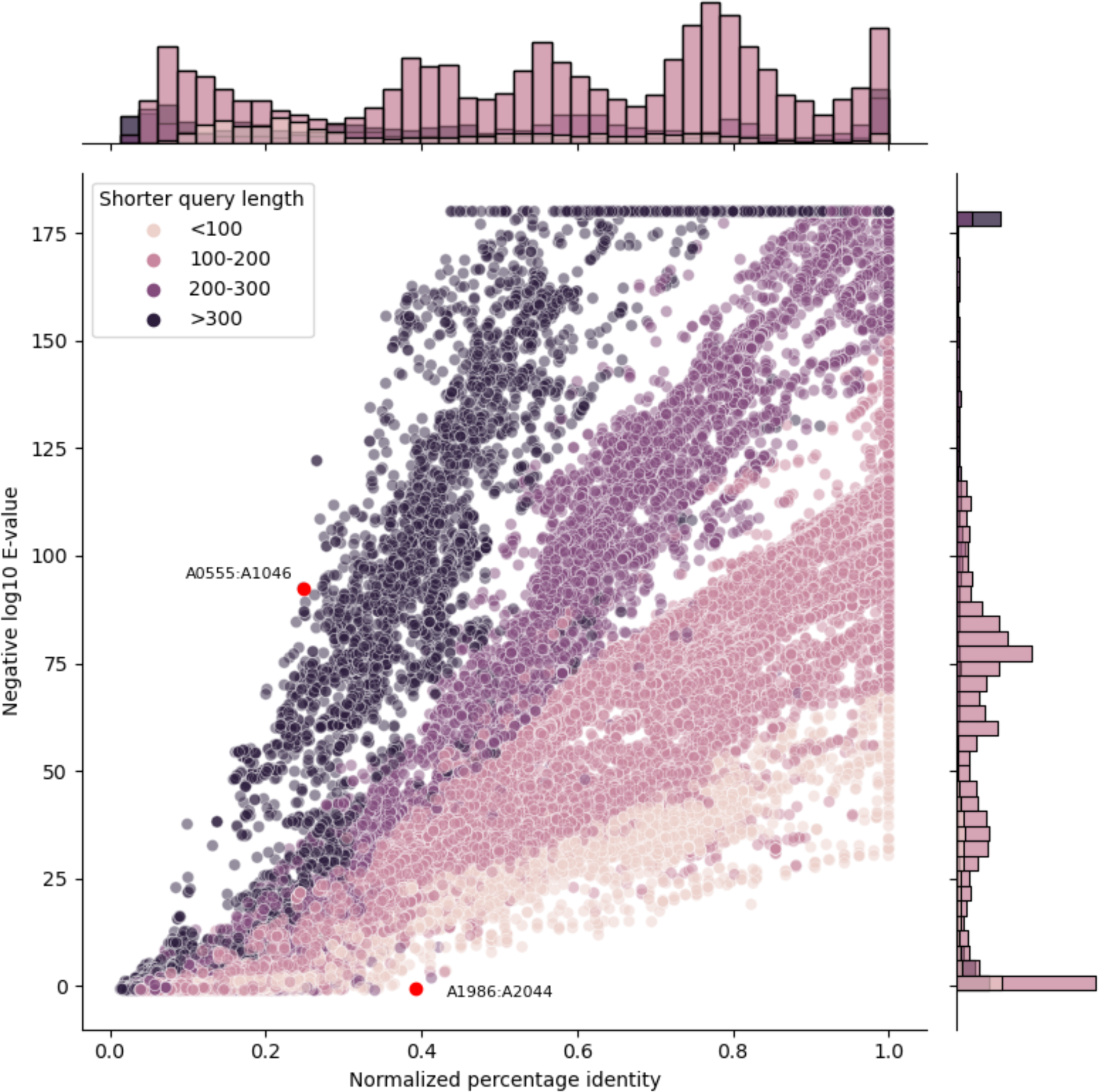
Comparison of E-value and normalized percentage identity of all-against-all BLAST alignment in the Allergen dataset 1. The E-value from BLAST was transformed to negative log E-value (-logE) for visualization purposes. Colors from pink to purple indicate lengths of query sequences. The upper histogram shows the count distribution of NPID values, and the distribution of -logE is shown in the right histogram. The red points represent two extreme alignment pairs where the E-value and percentage identity disagree.: A0555-A1046, and A1986-A2044.

### Benchmark of clustering

After investigating the sequence alignment, we performed clustering using ProtParts to link all the proteins that shared sequence similarity above several thresholds, using the similarity metrics compared before. The clustering performance using E-value or NPID was evaluated on the InterPro annotated Dataset 2 using the adjusted mutual information (AMI) metric, which represents the consistency between clusters and labels. When comparing the clustering with the optimal E-value threshold to the one using the best NPID, the clustering using E-values reached higher AMI than using NPID, regardless of which label was used (homologous superfamily, family, or domain) (Figure 2). In terms of protein families, the AMI was overall higher than the domain and the homologous superfamily when using an E-value of 1x10^-9^ (Figure 2A and 2B). These results aligned with the definition of family, i.e., proteins sharing a common evolutionary origin or ancestor. It indicated that proteins in the same family were likely to be more similar at the sequence level. On the contrary, the domain label required lower similarity thresholds which resulted in the highest AMI (Figure 2A and 2B). Overall, this suggested that the E-value was capable of finding more homologous sequences than the NPID.

In order to validate the E-value as a similarity metric, we further introduced a new annotated dataset (ASTRAL SCOPe), originating from PDB sequences that were classified into SCOPe domains. There are four labels for protein structures in the SCOPe database: class, fold, superfamily and family (from the most structure-based to the more evolutionary-based). We tested the clustering performance on this dataset comparing E-values and NPID (Figure 2C and 2D respectively). Again, the clustering using E-values compared to NPID showed a consistently higher AMI, suggesting it could be a better choice for protein clustering. Additionally, the clustering results with family labels, the most evolutionary-based, had the highest AMI under most thresholds both using E-values and NPID (Figure 2C, 2D); however, the maximum AMI of family was lower than that of superfamily using the optimal E-value (Figure 2C). The AMI of family and superfamily were followed by those of fold and class, which are more dependent on the conservation of protein structures but not necessarily on protein sequences. In addition, the clustering performance based on PID was evaluated to compare with -logE and NPID. However, the PID only had the highest AMI of 0.781 on the allergen dataset and 0.405 on the ASTRAL SCOPe dataset, which were lower than the AMI based on NPID and -logE.

**Figure 2.**
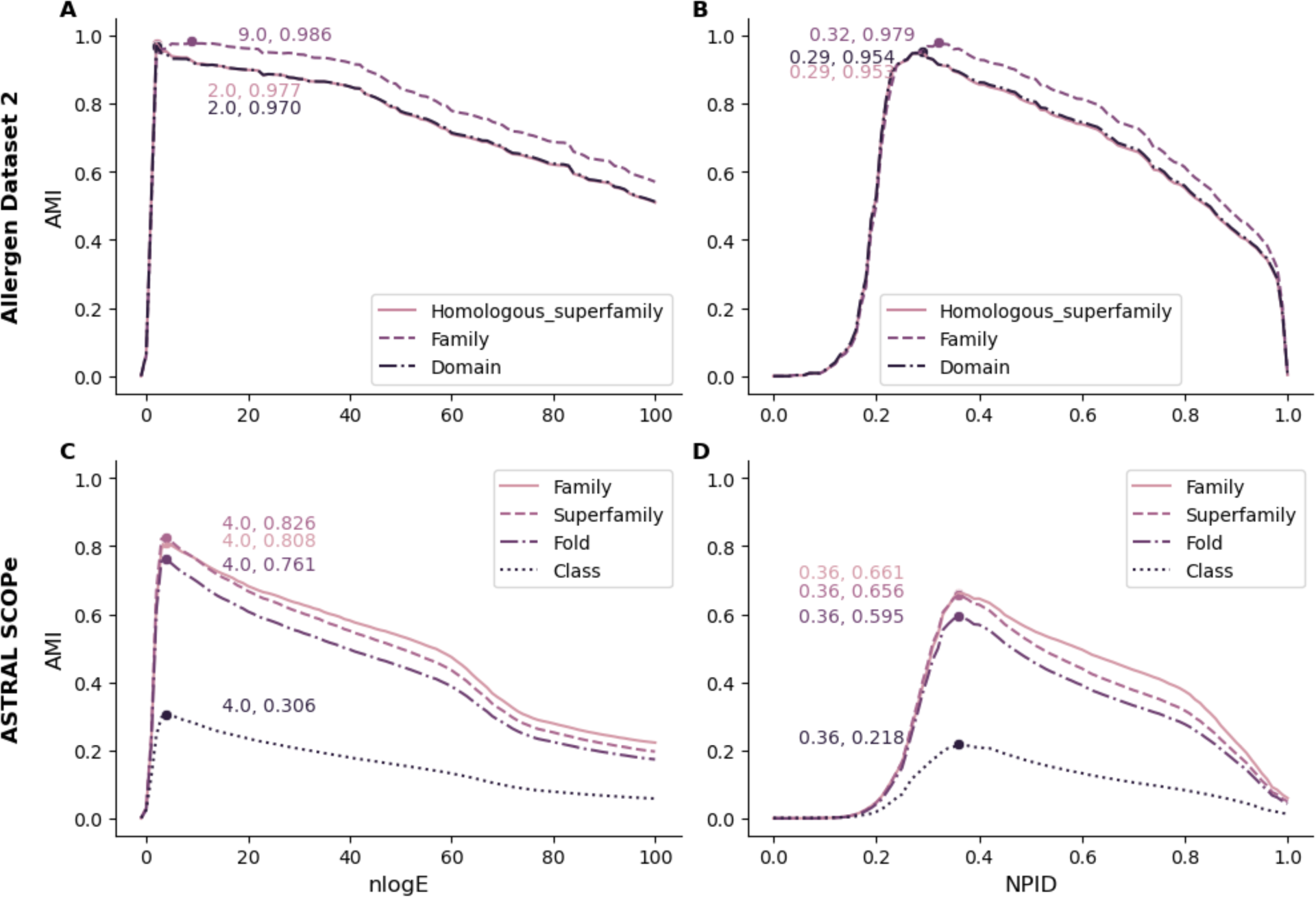
Comparison of clustering performance of ProtParts using E-values (A) vs NPID (B) on the Allergen dataset 2, or the ASTRAL SCOPe dataset (C, and D). The adjusted mutual information (AMI) was used to evaluate performance using different E-values or NPID thresholds. The E-value from BLAST was transformed to negative log E-value (-logE) for visualization purposes. AMI was calculated for homologous superfamily, family, and domain on the allergen dataset. AMI for family, superfamily, fold and class on the ASTRAL SCOPe dataset. The highest AMI comparing different thresholds is indicated with a dot. Both AMI performance values, along with the best threshold to obtain them are indicated on the plot with the same colour as the curves.

The composition of each cluster in the allergen dataset 1 was further analyzed based on the InterPro labels. When using the E-value of 1x10^-9^ as found to be the one with optimal AMI as clustering threshold, there were 215 clusters. As an example, several representative clusters corresponding to the most common seven allergen domains were illustrated (Figure 3). Superimposing the domain labels to the unsupervised clustering based on sequence similarity showed that most clusters contain a single domain or domain combination (Figure 3). Cluster 0 was the largest cluster containing 180 proteins, of which 177 were annotated with Bet v 1/Major latex domain and 3 were missing domain label association. Most other clusters at domain level exclusively included proteins with a single domain. For instance, the Cluster 5 (EF-hand domain) and the Cluster 12 (CAP domain) were separately grouped into uniform clusters as well. There were several clusters comprising a mix of single-domain and combined-domain proteins, while the combined-domain proteins had the exact same domain as the single-domain proteins, along with an extra domain, such as Cluster 9. In these cases, the majority of the proteins in the cluster contained two domains, and the minority contained only one of the domains. For instance, proteins with peptidase C1A and cathepsin propeptide inhibitor domain were the major sequences in Cluster 9, and the similarity of minor proteins came from the shared peptidase C1A domain. Although most proteins in the same domain were also similar in sequence, there was an exception that a domain had very distinct protein sequences. Proteins with the domain IPR016140 label were split into four different clusters because of the sequence homology, for example Cluster 3, 10, 13 and 19.

**Figure 3.**
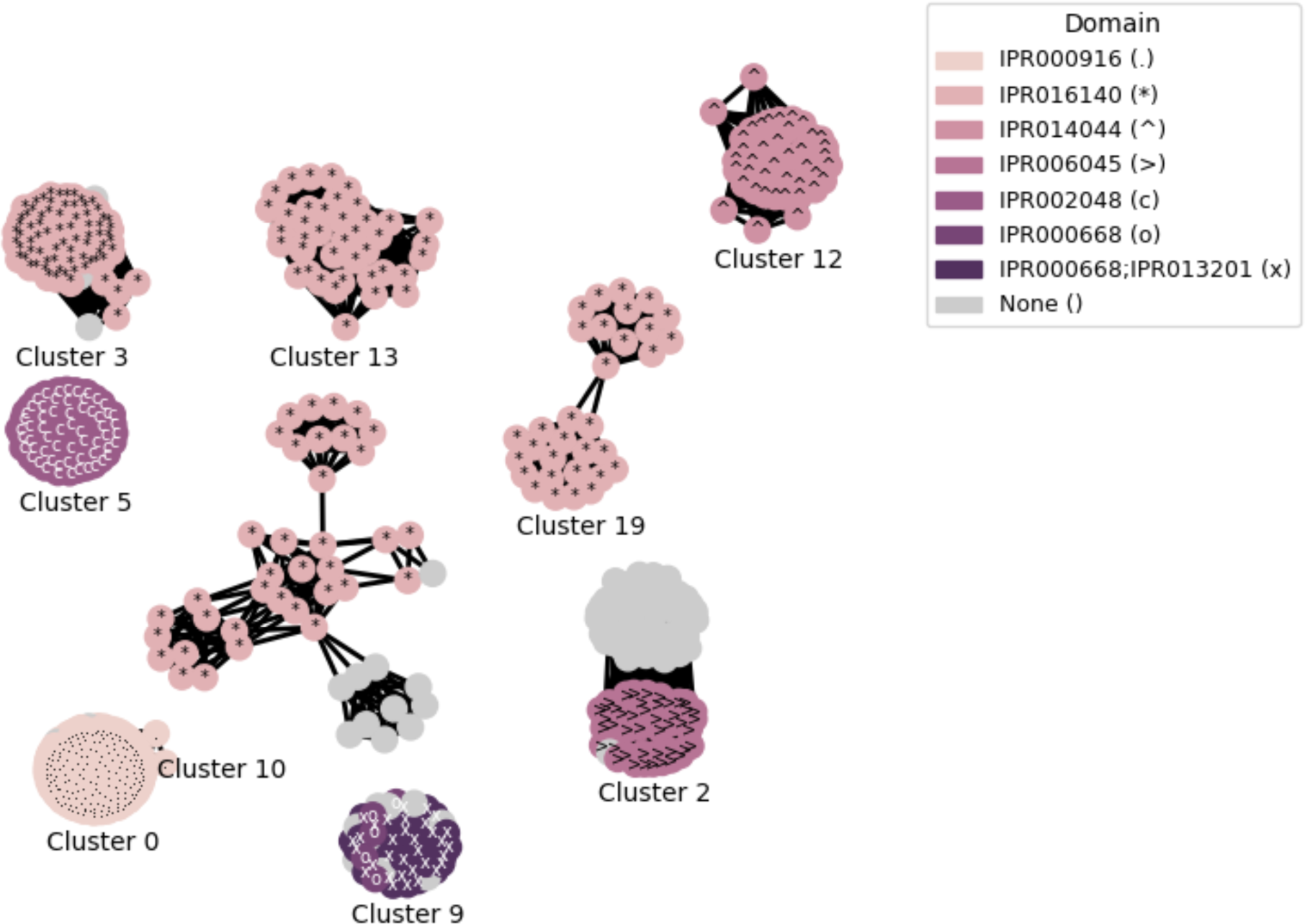
Examples of large clusters using ProtParts with E-value threshold of 1x10^-9^. Structural annotations of ‘domain’ labels from Dataset 2 are superimposed to the unsupervised clustering based on Dataset 1. There are seven common domains: Bet v I/Major latex protein (IPR000916), Bifunctional inhibitor/plant lipid transfer protein/seed storage helical domain (IPR016140), CAP domain (IPR014044), Cupin 1 (IPR006045), EF-hand domain (IPR002048), Peptidase C1A, papain C-terminal (IPR000668), Cathepsin propeptide inhibitor domain (I29) (IPR013201). One of them is the joint of two domains which suggests the protein contains both domains.

As the Cluster 0, 2, 3 and 5 exhibited homogeneities in terms of domain, the consistency was also maintained when they were divided at family, and superfamily level (Supplementary Figure S3 and S4). The clusters with a mix of single and combined domains, like Cluster 9, became more uniform clusters at family level. Nevertheless, proteins from the same domain could have different protein families. The Cluster 3, 10, 13 and 19 were the IPR016140, however, they were annotated with four different protein families: plant non-specific lipid-transfer protein/Par allergen, napin/2S seed storage protein/conglutin, gliadin/LMW glutenin, and cereal seed allergen respectively. These explained the AMI evaluation at the protein family label had the highest score. The same case applied to the Cluster 12 as well. Although the proteins in Cluster 12 belonged to the CAP domain, there were also three protein families within Cluster 12: cysteine-rich secretory protein-related, venom allergen 3, and the combination of these two families. Regarding the homologous superfamily, the clustering results provided more uniform clusters since the superfamily was higher than family or domain in protein structure hierarchy (Supplementary Figure S4).

Additionally, for comparison of clustering tools, we clustered these two datasets and compared the AMI performance across distinct tools: Hobohm1 (NPID), CD-HIT v4.6.8 (NPID), MMseqs2 v14-7e284 (PID+Evalue+query coverage), UCLUST v11.0 (PID), and ProtParts (NPID or E-value). Each tool generated clusters using a series of thresholds in either sequence identity or E-value (Supplementary Table S1). However, although most tools used the PID or NPID for clustering, these tools employed different approaches to measure the sequence identity. Therefore, for each tool, we exclusively selected and compared the cluster with the highest AMI selected from a range of thresholds (Figure 4). As a result, ProtParts using E-value outperformed the other tools based on sequence identity (CD-HIT, MMseqs2, UCLUST, Hobohm 1) for both datasets (Figure 4A, 4B). When using ProtParts based NPID, our tool outperformed CD-HIT/MMseq2 on the allergen dataset (Figure 4A); but not on the ASTRAL SCOPe dataset (Figure 4B). The comparable clustering performance of ProtParts (NPID) and CD-HIT/MMseqs2 on this dataset was probably derived from the different redundancy in these two datasets and different sequence identity metrics in these two clustering tools.

**Figure 4.**
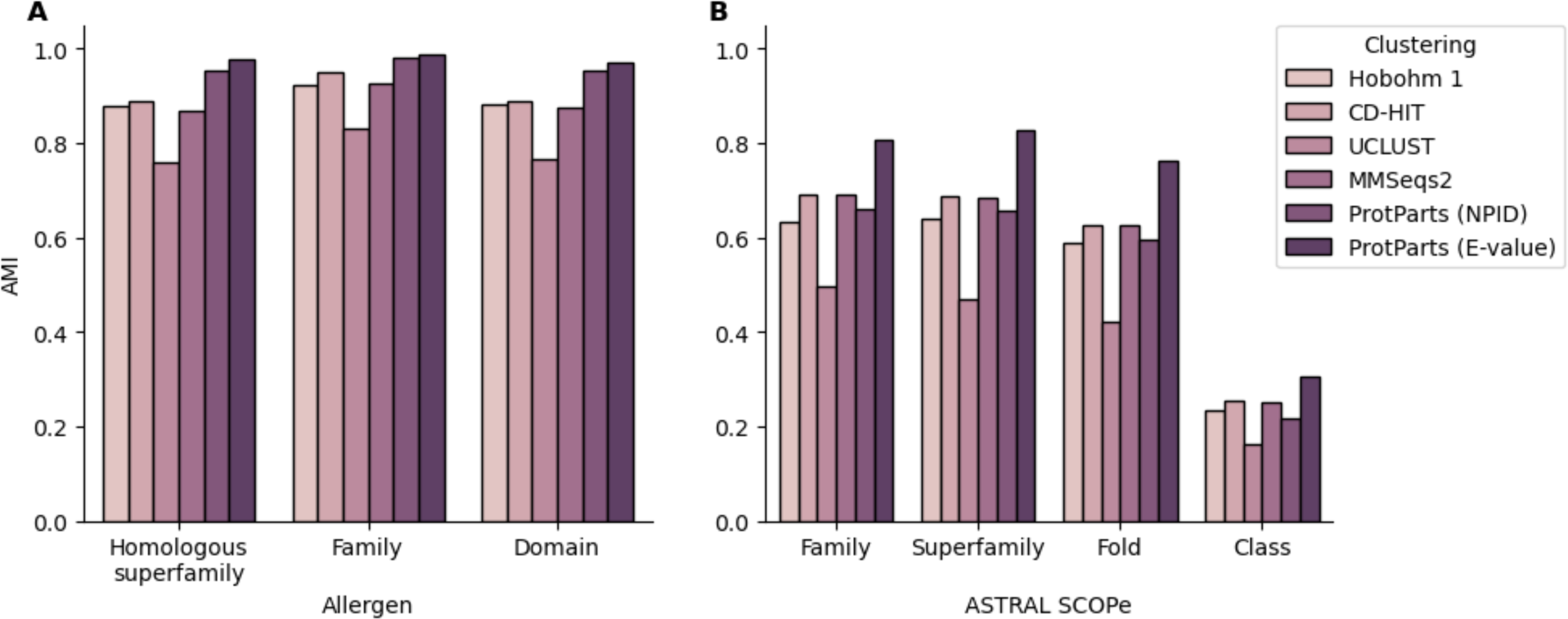
Adjusted mutual information (AMI) performance of clustering tools at the optimal threshold on structurally annotated protein dataset. The Hobohm 1 clustering used NPID as thresholds. CD-HIT, MMseqs2 and UCLUST were clustered based on their own PID. **A**. AMI on the allergen dataset with their optimal thresholds for homologous superfamily, family and domain B. AMI on the ASTRAL SCOPe dataset with their optimal thresholds for family, superfamily, fold and class.

### Machine learning performance

In this section, we selected three distinct clustering algorithms (ProtParts, Hobohm 1, and random) with the same similarity metric and threshold to test how the clustering affects machine learning training and evaluation performance. We firstly performed these three clustering approaches using E-value as a similarity metric and applied the same E-value threshold 1x10^-9^, obtained from the previous analysis, except for the random clustering that is agnostic to the E-value. We created the Hobohm 1-based clustering, utilizing an identical similarity measurement to ProtParts E-value of 1x10^-9^). Additionally, we created the random clustering by defining the number of clusters to 230 to approximately equal to the amount of clusters in both ProtParts and Hobohm 1, and to reduce the impact of the amount of clusters. Protein sequences were randomly allocated to these 230 clusters. As we concluded before comparing each method’s best threshold, the AMI of ProtParts showed a better agreement of clustering labels with structural label annotations than Hobohm 1 for the clustering generated using an E-value of 1x10^-9^ (Table1). In addition, both clustering approaches outperform the random clustering. Moreover, to comprehensively evaluate the performance of these clustering approaches, an unsupervised metric, the silhouette coefficient was assesed. Regardless of the labels, the silhouette coefficient measures the cohesion of a given cluster and the separation to other clusters. Since labels are not required for this assessment, the entire allergen dataset, Dataset 1, was evaluated. ProtParts also had the highest silhouette coefficient score, thereby indicating that ProtParts was able to reduce the similarities between partitions (Table 1). This superiority was consistent across all evaluation metrics indicating ProtParts had a more robust and accurate clustering capability than both the Hobohm 1 and random clustering methods.

**Table 1.**
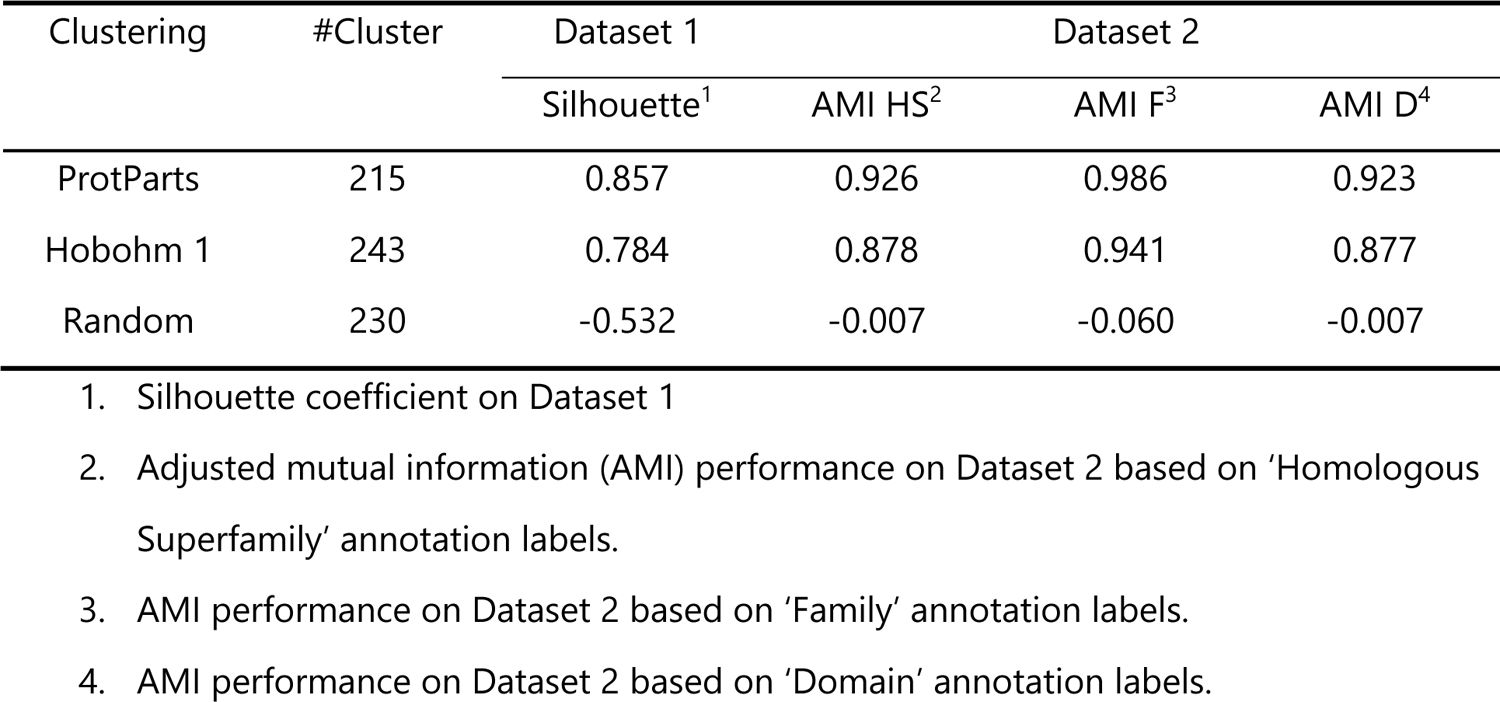
Evaluation performance of clustering methods on Dataset 1 and Dataset 2.

ProtParts connects all the proteins below a given threshold to generate partitions. However, this fact could lead to the formation of large clusters, sometimes exceeding the maximum capacity of one partition, affecting the following partitioning. To address this problem, we can remove the nodes which connect separated communities as a single cluster and then reduce the size of larger clusters. ProtParts has a function to remove the furthest node (which is measured by the ratio of the mean distance of this node to its neighbors over the mean distance of its neighbors to their neighbors) from the graph and re-clusters. This step is iteratively executed until the silhouette coefficient does not increase. By pruning the clusters, the silhouette coefficient on the allergen Dataset 1, as an example, had improved to 0.911 from 0.857 (Supplementary Figure S5). The pruning removed 28 nodes and created four more clusters. Compared with removing random nodes in the same graph, the optimal pruning, selecting the furthest one from the graph, led to higher silhouette coefficient (Supplementary Figure S5A). Additionally, the absolute silhouette coefficient between the optimal pruning and the node removal of baseline random clustering was increased as well (Supplementary Figure S5B, C). These results suggested that the cluster pruning generated better clusters with higher silhouette coefficient and reduced the size of large clusters consisting of different subgraphs.

In order to investigate the influence of clustering and partitioning on machine learning performance, we employed three predictors to train classification models on the full allergen dataset (Dataset 1). The allergen dataset was used to retrain three distinct models: NetAllergen, which employs a random forest algorithm; DeepAlgPro, which is based on a convolutional neural network architecture; and ProPythia, a neural network that incorporates descriptors. Meanwhile, a nested cross-validation was applied during the model training to prevent overfitting. In the cross-validation, the clusters generated in the preceding methods were randomly allocated to five partitions. As the cluster independence demonstrated by ProtParts, these partitions were likewise created to be independent, thereby preventing data leakage. To ensure independence in the nested cross-validation, the outer layer was composed of five folds, while the remaining four folds were utilized in the inner layer. The prediction of the outer layer test fold was an ensemble of predictions by four inner layer models. Concatenating five outer layer test folds, the final prediction was evaluated by AUC and AUC 0.1. In the random clustering, homologous data existed across different partitions (Supplementary Figure S6), therefore it was expected that random clustering always led to the highest performance in cross-validation, followed by Hobohm 1, and ProtParts (Figure 5A and B). ProtParts had the lowest AUC and AUC 0.1 as expected, because the partitions shared minimal similarity.

These models were further tested on a new evaluation allergen dataset (Dataset 3) to assess their predictive capabilities with data not previously encountered during training. The full evaluation dataset comprised a mix of sequences that were both homologous to the training dataset and distinct from them. First, we calculated the performance using a BLAST baseline model, in which each sequence in the evaluation dataset is searched against the allergen sequences in the training dataset, in this case the prediction value is the similarity metric. The baseline model prediction had an AUC of 0.941 and an AUC 0.1 of 0.868. This high AUC values suggested that sequence similarity to the train dataset was very high. To fully understand the influence of sequence similarity among the evaluation and training datasets on model performance, we sorted the evaluation dataset by high to low similarity to the training dataset, and removed one by one each of the proteins in the dataset. For each subset evaluation dataset, we evaluated model performance by AUC and AUC 0.1 (Figure 5C and 5D). These results showed that the model performance drastically decreased when similar sequences to the train dataset were removed. We can also observe how the different machine learning architectures tried here; random forest (NetAllergen), CNN(ProPythia), and deep neural network (DeepAlgPro) are more or less dependent on shared similarity across train and evaluation datasets and clustering methods. The random forest model showed to be less dependent on the clustering method used prior training. On the other hand, the deep model (DeepAlgPro), showed a big difference in performance when using random clustering to generate the training partitions and the evaluation dataset contained highly similar sequences to the training dataset. However, this performance drastically drops until is lower than the clustering done with ProtParts when those highly similar sequences are removed from the dataset both in AUC and AUC 0.1 (Figure 5C and 5D, middle panels). These results together showed that the deep model was more sensitive to clustering and proper partitions on the training dataset and might show inflated performances on cross-validation or homologous dataset while performing very poorly on unseen data. Therefore, seems extremely relevant to user proper clustering and partition for models using protein sequences.

**Figure 5.**
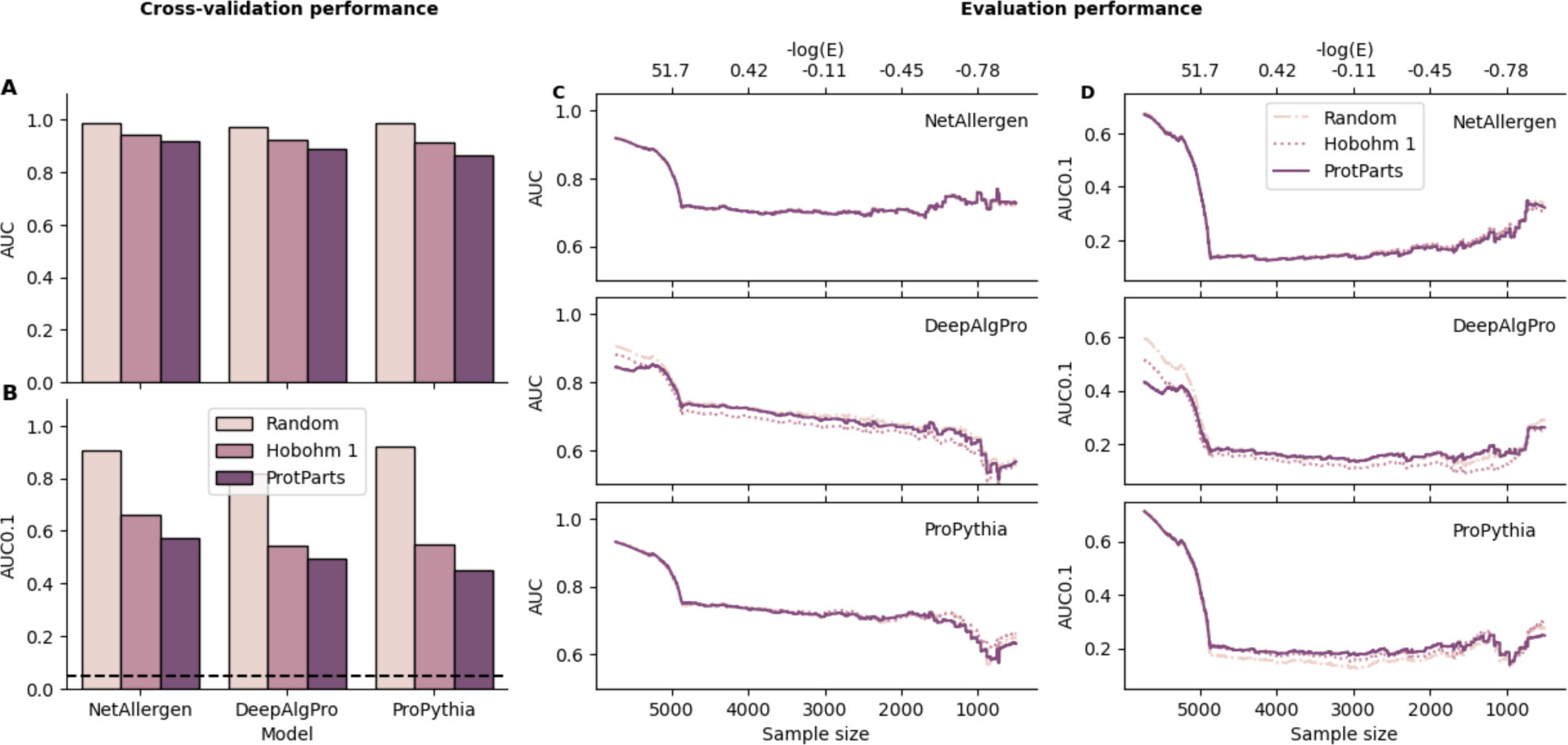
Model performance on allergen dataset. **A, B**. AUC, AUC 0.1 on the cross-validation training dataset. The dashed line is the AUC 0.1 of a random predictor, which is approximately 0.05. **C**. AUC machine learning performance of three different clusterings for NetAllergen, DeepAlgPro and ProPythia on evaluation dataset. **D**. AUC 0.1 performance on the evaluation dataset. **C, D.** AUC, AUC 0.1 on the full evaluation dataset. **E, F**. AUC, AUC 0.1 on the independent evaluation dataset (a subset of the full evaluation dataset, with less similarities to training dataset). The dashed line in C and D is the AUC 0.1 for a random predictor.

### Web server

ProtParts has been deployed as a user-friendly web server at https://services.healthtech.dtu.dk/services/ProtParts-1.0/ (Supplementary Figure S7). ProtParts deals with a protein FASTA sequence file, provides multiple choices for clustering thresholds, and generates diverse output formats to show the best possible clustering on sequence similarity. We enable two optional functions on the ProtParts web server. First, users are able to reduce sequence redundancy of their dataset using the Hobohm 1 algorithm with a user defined E-value threshold. The other function is ‘Pruning to improve clustering performance’. This option will remove proteins from the dataset that are bridge nodes, or connect different dense subgraphs, to improve clustering performance assessed on the Silhouette coefficient.

## DISCUSSION

The issue of overfitting in machine learning is a critical concern that demands careful consideration. In bioinformatics, the presence of similar biological sequences in training and test dataset can often lead to overfitting. In this study, we proposed a protein sequence clustering and partitioning tool, ProtParts, which utilized E-value as sequence similarity measurement and employed graph approach to generate independent clusters. By applying ProtParts on different types of datasets, sequences in the same cluster were proved to share more accurate protein structural information than using percentage identity as thresholds. Moreover, the graph of ProtParts ensured that no data or information leakage occurred among clusters and partitions and achieved a better clustering performance. Compared to random clustering in machine learning, the independence of ProtParts partitions effectively avoided overfitting and overestimated predictive performance in test datasets.

In our comprehensive analysis of E-values and NPID derived from BLAST results, we observed that the E-value carried more biological information and could be regarded as significant sequence similarity. It was also proved to be more beneficial for protein clustering by assessing the AMI. The sequence alignment comparison revealed that the E-value was effective to discover longer and more complex homologous sequences. The effectiveness can be attributed to the fact that the E-value considers evolutionary substitution not only the identically aligned amino acids, which are likely crucial in finding the differences in clustering (20). In contrast, NPID, which measures the normalized percentage of identical matches by the shorter sequence length, was more applicable to find shorter and simpler similar sequences. Additionally, by comparing the AMI of clusterings based on the NPID and PID, the NPID was able to provide a better discrimination for protein sequences. Nevertheless, when we evaluated the clustering outcomes using different metrics, such as the silhouette coefficient and the adjusted mutual information, the E-value consistently demonstrated superior clustering performance. This was indicative of its capacity of protein clustering, not only achieving a good separation in terms of sequence similarity, but also ensuring that different clusters were biologically meaningful.

In the case of employing E-value as a threshold, the evaluation benchmark of clustering methods demonstrated that ProtParts ensured the independence among clusters. Hobohm 1 clustering only compares query sequences with the representative sequence of each cluster, which can efficiently cluster sequences. However, the similarities among other member sequences from different clusters were still unknown. This would most likely lead to the existence of similar sequences in many different clusters, thereby leading to data leakage in machine learning partitions. Unlike the Hohobm 1 clustering which only compared to representative protein sequences, the ProtParts assessed all-against-all sequence similarity. It was precisely the all-against-all comparison that resulted in the highest silhouette coefficient of ProtParts. Moreover, the AMI of ProtParts was the highest at all different structural labels, homologous superfamily, family and domain. ProtParts outperformed other commonly used tools (CD-HIT, MMseq2 and UCLUST) in the comparison of AMI as well.

Upon comparing clustering results in machine learning, it was evident that training a model on partitions from random clustering tends to yield inflated performance, even if the evaluation dataset had the similar distribution to the cross-validation dataset. As a result, when there was data leakage among partitions, the model performance on the cross-validation may not accurately reflect their actual predictive efficacy. However, the models trained on partitions from ProtParts were able to maintain a close performance on the same evaluation dataset. It indicated that independent partitions in machine learning could largely avoid overestimation of test data prediction. On the filtered evaluation dataset, the true independent data, the models trained with ProtParts achieved better performance or at least similar performance, indicative of less overfitting. They demonstrated an ability to accurately predict data that was not previously encountered. Compared to neural network based machine learning algorithms, the clustering had less influence on the training and evaluation performance for the random forest model. This was probably because the random forest included bootstrapping of data and features in the initial algorithm, thereby it was less sensitive to different clustering or partitioning.

Data leakage stands as a critical issue in machine learning, potentially compromising the reliability of model prediction. That data leakage in different forms leads to exaggerated performance has become a consensus in the scientific community (34, 35). In our research, to solve the data leakage issue, we have substantiated that a proper clustering and partitioning of data contributes to achieve more generalized and robust results, especially in deep learning. Intricate networks and large amounts of parameters make them prone to be overfitted. Additionally, the selection of sequence similarity metrics is basically the percentage identity. We utilized the E-value for clustering instead of the most frequently used percentage identity. It validated that the percentage identity is less informative than the E-value. The application of E-value improved the clustering performance. By implementing the ProtParts, we can discover proteins with shared properties, as well as ensure the independence of data across datasets in machine learning to avoid overfitting.

## DATA AVAILABILITY

The ProtParts web server, terminal program and relevant datasets are available at: https://services.healthtech.dtu.dk/services/ProtParts-1.0/

## ACKNOWLEDGEMENTS

We acknowledge Morten Nielsen and Henrik Nielsen for their useful comments on the paper.

