## Supplemental Figure for "ProtParts, an automated web server for clustering and partitioning protein dataset"

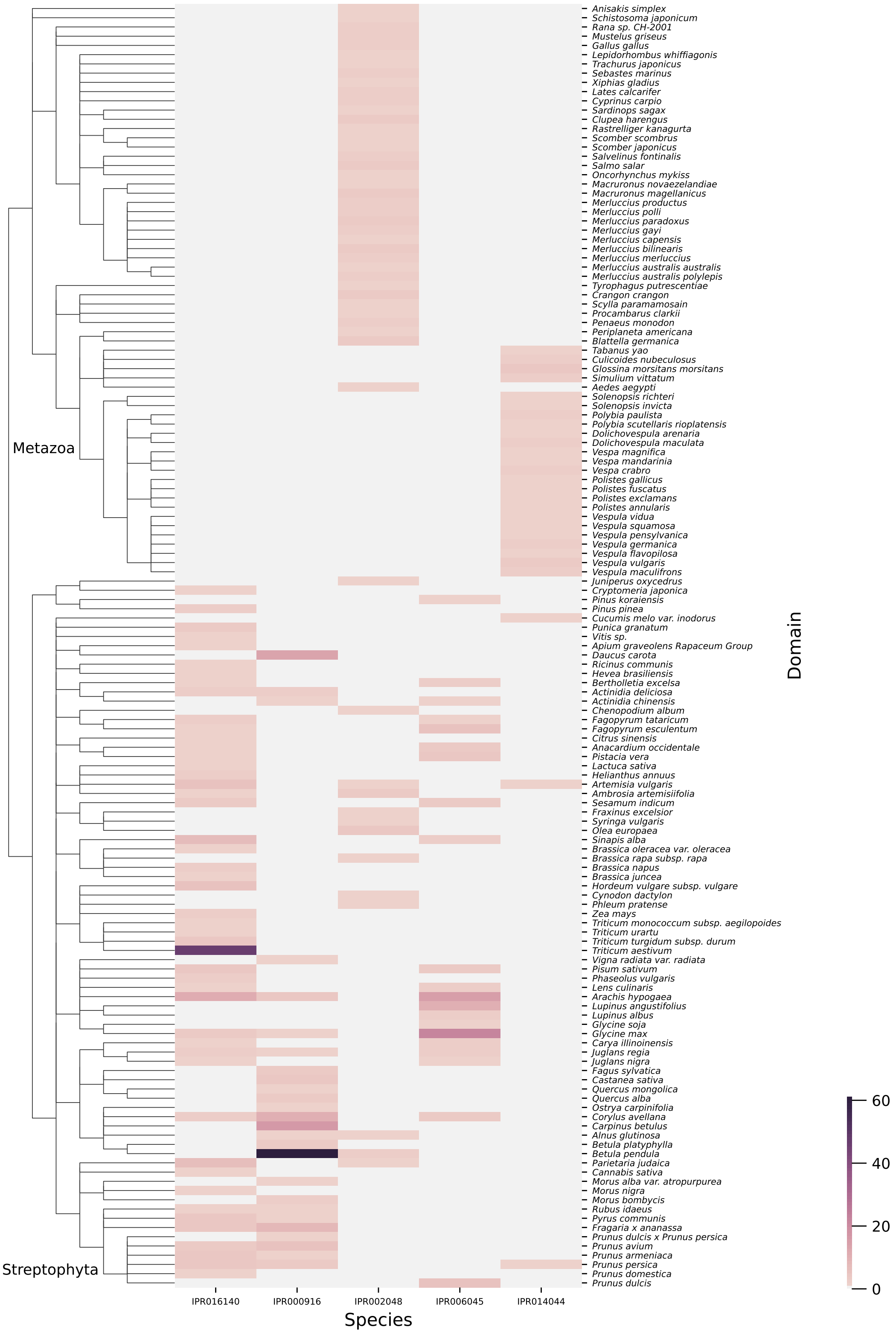


**Supplementary Figure S1.** Occurrences distribution of protein domains that cover more than 20 species. Species in the heatmap are rearranged by their evolutionary relationship from the NCBI taxonomy database. The InterPro IDs represent these domains: Bifunctional inhibitor/plant lipid transfer protein/seed storage helical domain (IPR016140), Bet v I/Major latex protein (IPR000916), EF-hand domain (IPR002048), Cupin 1 (IPR006045), CAP domain (IPR014044).

**A**

```
>A2130-677_1
Length=454

 Score = 253 bits (645),  Expect = 3e-77, Method: Compositional matrix adjust.
 Identities = 144/404 (36%), Positives = 257/404 (64%), Gaps = 26/404 (6%)

Query  375  RHNPYYFHSQ-GLRSRHESGEGEVKYLERFTERTELLRGIENYRVVILEANPNTFVLPYH  433
            ++NP+YF+S     +   +  G ++ L+RF +R++ ++ +ENYRVV L + PNT +LP+H
Sbjct  43   KNNPFYFNSDRWFHTLFRNQFGHLRVLQRFDQRSKQMQNLENYRVVELMSKPNTLLLPHH  102

Query  434  KDAESVIVVTRGRATLTFVSQERRESFNLEYGDVIRVPAGATEYVINQDSNERLEMVKLL  493
             DA+ ++VV  GRA LT V+ + R+S  LE G   ++PAG T +++N D NE L ++KL
Sbjct  103  ADADFLLVVLNGRAVLTLVNPDGRDSNILEQGHAQKIPAGTTFFLVNPDDNENLRIIKLA  162

Query  494  QPVNNPGQFREYYAAGAQSTESYLRVFSNDILVAALNTPRDRLERFF----------DQQ  543
             PVNNP +F++++ +  ++ +SYL+ FS +IL A+ ++    + R             Q+
Sbjct  163  VPVNNPHRFQDFFLSSTEAQQSYLQGFSKNILEASFDSDIKEISRVLFGEEGQQQQQGQE  222

Query  544  EQREGVIIRASQEKLRALSQHAMSAGQRPWGRRSSGGPISLKSQRSSYSNQFGQFFEACP  603
             Q+EGVI+   +E++R L++HA S+ ++     S   P +L++Q+  YSN+ G++FE  P
Sbjct  223  SQQEGVIVELKREQIRELTKHAKSSSKK--SLSSEDQPFNLRNQKPIYSNKLGRWFEITP  280

Query  604  EEHRQLQEMDVLVNYAEIKRGAMMVPHYNSKATVVVYVVEGTGRFEMACPHDVSSQSYEY  663
            E++ QL+++D+ +   ++K G++++PHYNSKA V++ + EG    E+
Sbjct  281  EKNPQLRDLDMFIRSVDMKEGSLLLPHYNSKAIVILVINEGKANIELVG----------Q  330

Query  664  KGRREQEEEESSTGQFQKVTARLARGDIFVIPAGHPIAITASQNENLRLVGFGINGKNNQ  723
            + +++Q+EE+  + + Q+  A L+  D+F+IPA +P+AI A+   NL    FGIN +NNQ
Sbjct  331  REQQKQQEEQEESWEVQRYRAELSEDDVFIIPATYPVAINATS--NLNFFAFGINAENNQ  388

Query  724  RNFLAG-QNNIINQLEREAKELSFNMPREEIEEIFERQVESYFV  766
            RNFLAG ++N+I+++  E  +++F    E+++++ ++Q ES FV
Sbjct  389  RNFLAGEKDNVISEIPTEVLDVTFPASGEKVKKLIKKQSESQFV  432
```

**B**

```
>A2044-653_1
Length=244

 Score = 24.3 bits (51),  Expect = 4.6, Method: Compositional matrix adjust.
 Identities = 96/248 (39%), Positives = 124/248 (50%), Gaps = 21/248 (8%)

Query  57   FPPQQPYPQQQPFPSQQPYMQLQPFPQPQLPYPQPQLPYPQPQPFRPQQSYPQPQPQYSQ  116
              PQQ +  Q   P  Q   Q  P PQ  +    P  P   P   +P  +
Sbjct  7    LQPQQSFLWQSQQPFLQQPQQPSPQPQQVVQIISPATPTTIPSAGKPTSA----------  56

Query  117  PQQPISQQQQQQQQQQQQQQQILQQILQQQLIPCRDVVLQQHS-IAH----GSSQVLQQS  171
               P  QQQQQ QQ  QQQ  ++Q  + QQL PC+  + QQ S +A       SQ+LQQS
Sbjct  57   ---PFPQQQQQHQQLAQQQIPVVQPSILQQLNPCKVFLQQQCSPVAMPQRLARSQMLQQS  113

Query  172  TYQLVQQFCCQQLWQIPEQSRCQAIHNVVHAIILHQQQQQQQQQQQQQQQPLSQVCFQQS  231
            +  ++QQ CCQQL QIP+QSR QAI  ++++IIL +QQQ Q   Q QQQQP
Sbjct  114  SCHVMQQQCCQQLPQIPQQSRYQAIRAIIYSIILQEQQQVQGSIQSQQQQPQQLGQCVSQ  173

Query  232  QQQYPSGQGSFQPSQQNPQAQGSVQPQQLPQFEEIRNLALETLPAMCNVYIPPY--CTIA  289
             QQ    Q   QP QQ        +QP Q+ Q E + ++AL  LP MC+V +P Y   T
Sbjct  174  PQQQSQQQLGQQPQQQQLAQGTFLQPHQIAQLEVMTSIALRILPTMCSVNVPLYRTTTSV  233

Query  290  PVGIFGTN  297
            P G+ GT
Sbjct  234  PFGV-GTG  240
```

**Supplementary Figure S2.** Sequence alignments of **A**. A0555 and A2130 with a NPID of 18%, **B**. A1980 and A2044 with a NPID of 32%.

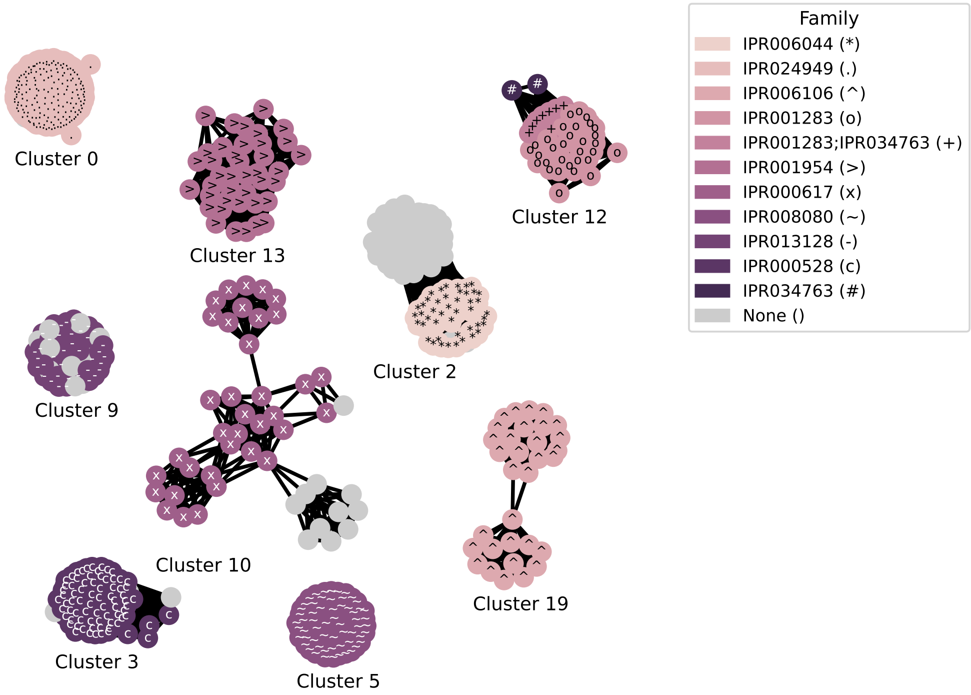


**Supplementary Figure S3.** Demonstration of partial clusters in a graph with family annotation. It displays the following families: 11-S seed storage protein, plant (IPR006044), Bet v I type allergen (IPR024949), Cereal seed allergen/grain softness/trypsin and alpha-amylase inhibitor (IPR006106), Cysteine-rich secretory protein-related (IPR001283), Gliadin/LMW glutenin (IPR001954), Napin/ 2S seed storage protein/Conglutin (IPR000617), Parvalbumin (IPR008080), Peptidase C1A (IPR013128), Plant non-specific lipid-transfer protein/Par allergen (IPR000528), Venom allergen 3, insect (IPR034763).

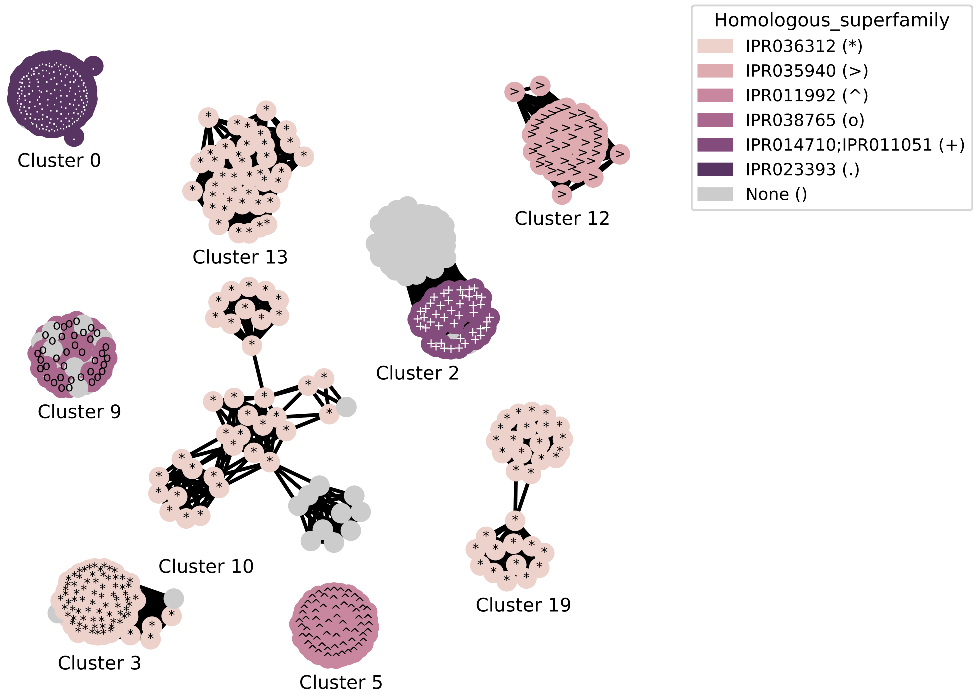


**Supplementary Figure S4.** Demonstration of partial clusters in a graph with homologous superfamily annotation. Seven unique homologous superfamily are shown: Bifunctional inhibitor/plant lipid transfer protein/seed storage helical domain superfamily (IPR036312), CAP superfamily (IPR035940), EF-hand domain pair (IPR011992), Papain-like cysteine peptidase superfamily (IPR038765), RmlC-like jelly roll fold (IPR014710), RmlC-like cupin domain superfamily (IPR011051), START-like domain superfamily (IPR023393).

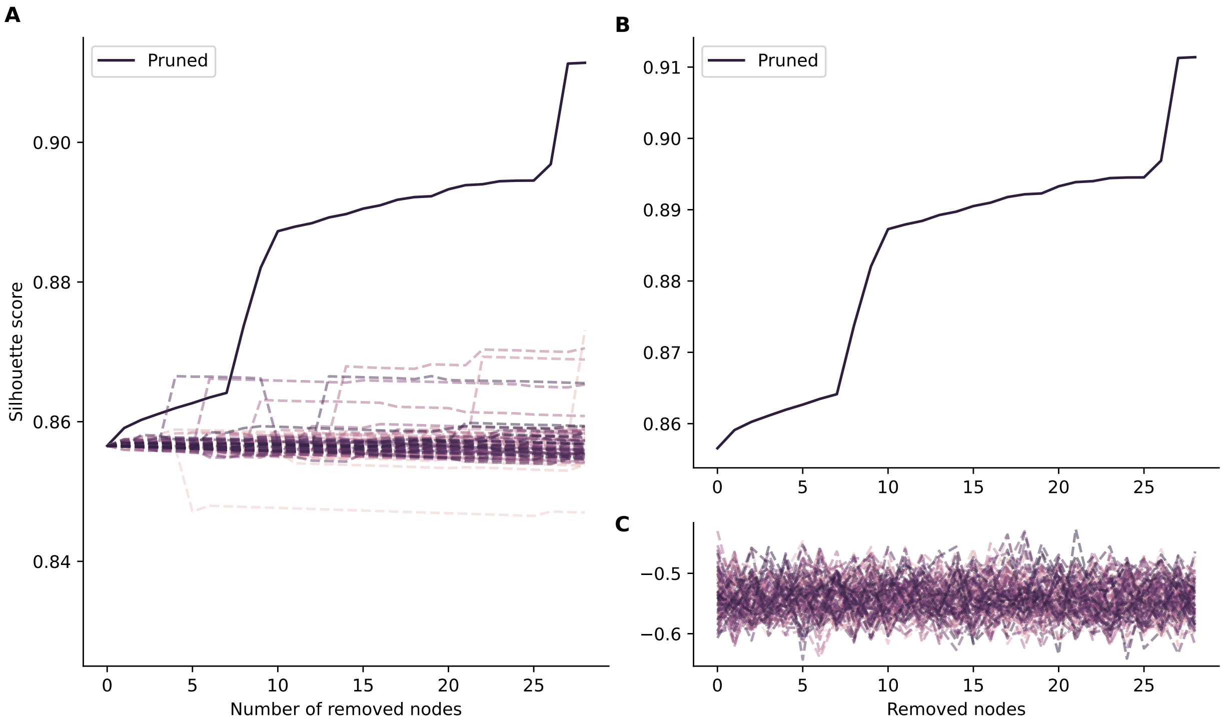


**Supplementary Figure S5.** Comparison of optimal pruned clusters with two different randomly pruned clusters. **A**. The solid line represents the optimal pruning and the dashed lines are for 100 pruning random samples from the same ProtParts clustering. Silhouette coefficient of removing identical samples from **B** ProtParts clustering and 100 **C** random clustering.

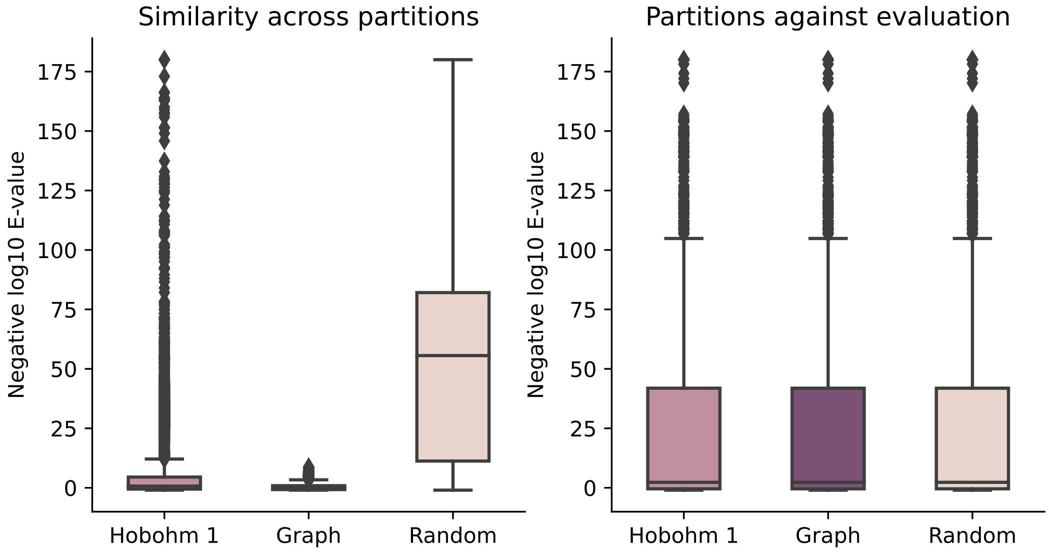


**Supplementary Figure S6.** Sequence similarity distribution (-logE) for different clustering methods **A**. across training partitions in the cross-validation. **B**. between training partitions and evaluation dataset.

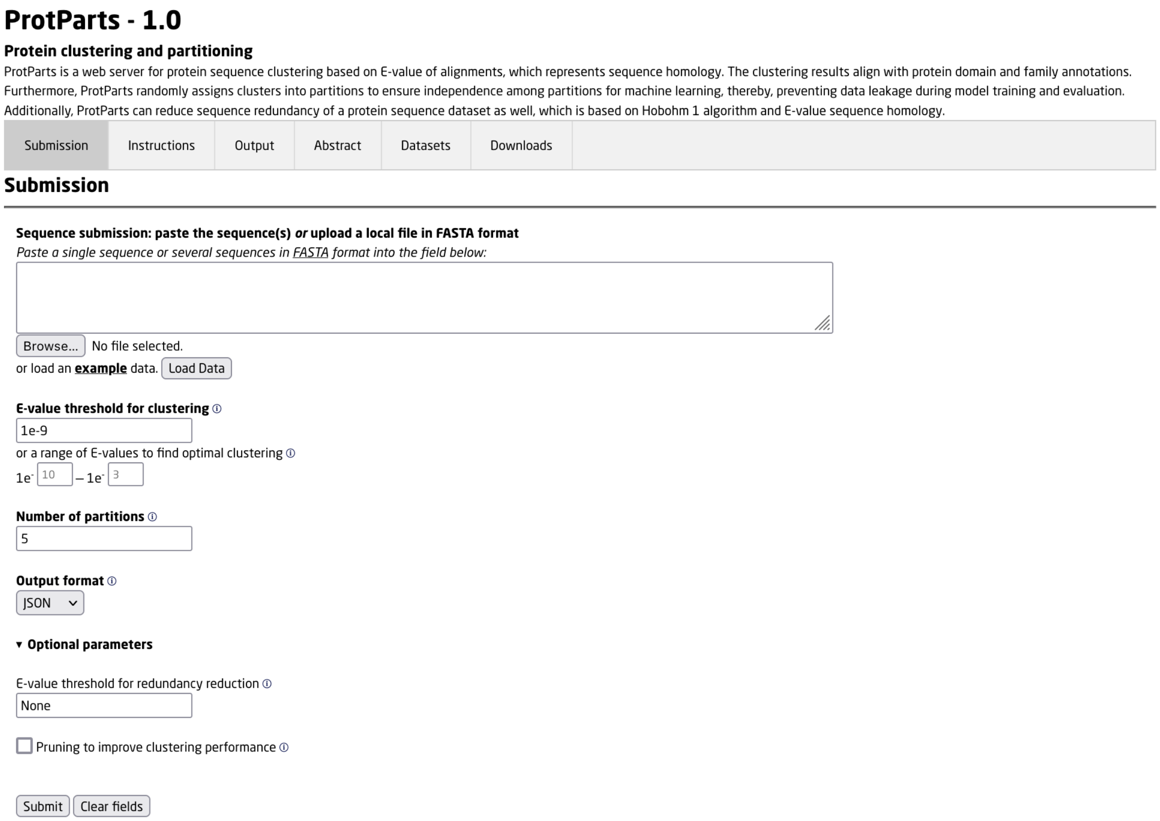


**Supplementary Figure S7.** ProtParts web server

**Supplementary Table S1.** Adjusted mutual information (AMI) of clusterings on allergen (Dataset 2) and ASTRAL SCOPe dataset at series of thresholds.

| Clustering | Threshold | Allergen | | | | ASTRAL SCOPe | | | | |
|---|---|---|---|---|---|---|---|---|---|---|
| | | #Cluster | AMI HS[1] | AMI F[2] | AMI D[3] | #Cluster | AMI Fa[4] | AMI S[5] | AMI Fo[6] | AMI C[7] |
| | 0.1 | 194 | 0.772 | 0.792 | 0.768 | 5029 | 0.508 | 0.514 | 0.471 | 0.175 |
| | **0.2** | **232** | **0.878** | **0.923** | **0.88** | **7148** | **0.632** | **0.641** | **0.588** | **0.235** |
| | 0.3 | 281 | 0.843 | 0.908 | 0.847 | 10481 | 0.607 | 0.609 | 0.556 | 0.225 |
| | 0.4 | 353 | 0.781 | 0.852 | 0.786 | 14794 | 0.531 | 0.517 | 0.469 | 0.184 |
| Hobohm 1 | 0.5 | 443 | 0.725 | 0.797 | 0.731 | 18591 | 0.468 | 0.437 | 0.393 | 0.147 |
| | 0.6 | 528 | 0.641 | 0.711 | 0.647 | 21713 | 0.412 | 0.371 | 0.332 | 0.119 |
| | 0.7 | 621 | 0.593 | 0.656 | 0.599 | 24365 | 0.337 | 0.298 | 0.266 | 0.094 |
| | 0.8 | 776 | 0.483 | 0.538 | 0.488 | 27010 | 0.244 | 0.21 | 0.188 | 0.07 |
| | 0.9 | 999 | 0.381 | 0.427 | 0.385 | 30583 | 0.118 | 0.101 | 0.09 | 0.037 |
| | **0.1** | **250** | **0.887** | **0.951** | **0.887** | **8943** | **0.691** | **0.686** | **0.626** | **0.255** |
| | 0.2 | 256 | 0.882 | 0.945 | 0.882 | 9171 | 0.676 | 0.672 | 0.614 | 0.25 |
| | 0.3 | 306 | 0.831 | 0.904 | 0.836 | 11295 | 0.611 | 0.606 | 0.552 | 0.222 |
| | 0.4 | 328 | 0.799 | 0.868 | 0.805 | 14376 | 0.531 | 0.514 | 0.465 | 0.178 |
| CD-HIT | 0.5 | 411 | 0.737 | 0.809 | 0.744 | 18467 | 0.47 | 0.438 | 0.394 | 0.148 |
| | 0.6 | 490 | 0.652 | 0.721 | 0.658 | 21677 | 0.403 | 0.363 | 0.325 | 0.118 |
| | 0.7 | 602 | 0.591 | 0.654 | 0.597 | 24397 | 0.322 | 0.287 | 0.257 | 0.093 |
| | 0.8 | 747 | 0.49 | 0.546 | 0.495 | 27017 | 0.237 | 0.205 | 0.184 | 0.069 |
| | 0.9 | 973 | 0.389 | 0.435 | 0.393 | 30647 | 0.115 | 0.099 | 0.088 | 0.037 |
| | 0.1 | 339 | 0.753 | 0.821 | 0.756 | 15164 | 0.478 | 0.454 | 0.41 | 0.157 |
| | **0.2** | **359** | **0.76** | **0.829** | **0.764** | **16451** | 0.493 | 0.468 | 0.422 | 0.161 |
| | **0.3** | 367 | 0.756 | 0.828 | 0.76 | 17158 | **0.496** | **0.468** | **0.422** | **0.161** |
| | 0.4 | 379 | 0.754 | 0.827 | 0.76 | 17532 | 0.492 | 0.462 | 0.416 | 0.158 |
| UCLUST | 0.5 | 433 | 0.71 | 0.781 | 0.717 | 18973 | 0.461 | 0.426 | 0.383 | 0.144 |
| | 0.6 | 506 | 0.667 | 0.736 | 0.673 | 21917 | 0.386 | 0.347 | 0.311 | 0.115 |
| | 0.7 | 616 | 0.566 | 0.627 | 0.572 | 24538 | 0.307 | 0.271 | 0.243 | 0.091 |
| | 0.8 | 770 | 0.48 | 0.533 | 0.485 | 27239 | 0.225 | 0.195 | 0.174 | 0.067 |
| | 0.9 | 1000 | 0.381 | 0.428 | 0.384 | 30881 | 0.107 | 0.091 | 0.082 | 0.035 |
| | **0.1** | 327 | 0.869 | 0.925 | 0.874 | **9490** | **0.69** | **0.683** | **0.625** | **0.251** |
| | **0.2** | **327** | **0.869** | **0.925** | **0.874** | 9495 | 0.69 | 0.683 | 0.624 | 0.251 |
| | 0.3 | 349 | 0.854 | 0.911 | 0.86 | 11036 | 0.641 | 0.63 | 0.575 | 0.23 |
| | 0.4 | 397 | 0.815 | 0.876 | 0.823 | 14570 | 0.569 | 0.546 | 0.494 | 0.191 |
| MMseqs2* | 0.5 | 473 | 0.741 | 0.802 | 0.749 | 18267 | 0.497 | 0.459 | 0.413 | 0.153 |

Continued on next page

**Continued**

| Clustering | Threshold | Allergen | | | | ASTRAL SCOPe | | | | |
|---|---|---|---|---|---|---|---|---|---|---|
| | | #Cluster | AMI HS[1] | AMI F[2] | AMI D[3] | #Cluster | AMI Fa[4] | AMI S[5] | AMI Fo[6] | AMI C[7] |
| | 0.6 | 547 | 0.693 | 0.752 | 0.701 | 21660 | 0.404 | 0.36 | 0.323 | 0.119 |
| | 0.7 | 642 | 0.596 | 0.652 | 0.603 | 24434 | 0.328 | 0.286 | 0.256 | 0.093 |
| | 0.8 | 782 | 0.497 | 0.545 | 0.503 | 27119 | 0.24 | 0.206 | 0.184 | 0.069 |
| | 0.9 | 1013 | 0.386 | 0.427 | 0.391 | 30795 | 0.115 | 0.098 | 0.088 | 0.036 |
| ProtParts (PID) | 0.1 | 28 | 0.015 | 0.015 | 0.018 | 32 | 0.001 | 0.002 | 0.001 | 0.002 |
| | 0.2 | 128 | 0.525 | 0.502 | 0.528 | 1401 | 0.049 | 0.048 | 0.043 | 0.020 |
| | **0.3** | **215** | **0.934** | **0.969** | **0.935** | 6750 | 0.446 | 0.459 | 0.419 | 0.160 |
| | **0.4** | 290 | 0.857 | 0.929 | 0.863 | **12706** | **0.645** | **0.628** | **0.568** | **0.209** |
| | 0.5 | 368 | 0.802 | 0.875 | 0.808 | 17334 | 0.555 | 0.517 | 0.464 | 0.167 |
| | 0.6 | 444 | 0.740 | 0.814 | 0.746 | 20847 | 0.496 | 0.440 | 0.391 | 0.133 |
| | 0.7 | 550 | 0.662 | 0.731 | 0.669 | 23695 | 0.437 | 0.377 | 0.332 | 0.106 |
| | 0.8 | 684 | 0.553 | 0.614 | 0.559 | 26174 | 0.374 | 0.317 | 0.277 | 0.083 |
| | 0.9 | 926 | 0.420 | 0.470 | 0.424 | 29340 | 0.227 | 0.193 | 0.169 | 0.052 |
| ProtParts (E-value) | 1E-01 | 129 | 0.520 | 0.488 | 0.514 | 3520 | 0.225 | 0.246 | 0.218 | 0.122 |
| | 1E-02 | 165 | 0.977 | 0.932 | 0.970 | 5172 | 0.629 | 0.669 | 0.609 | 0.245 |
| | 1E-03 | 184 | 0.954 | 0.960 | 0.949 | 5804 | 0.793 | 0.822 | 0.758 | 0.298 |
| | **1E-04** | 189 | 0.953 | 0.962 | 0.946 | **6287** | **0.807** | **0.824** | **0.759** | **0.306** |
| | 1E-05 | 195 | 0.939 | 0.976 | 0.932 | 6635 | 0.800 | 0.810 | 0.746 | 0.302 |
| | 1E-06 | 202 | 0.937 | 0.975 | 0.932 | 6975 | 0.791 | 0.798 | 0.734 | 0.296 |
| | 1E-07 | 207 | 0.936 | 0.976 | 0.932 | 7316 | 0.781 | 0.785 | 0.721 | 0.290 |
| | 1E-08 | 211 | 0.936 | 0.976 | 0.932 | 7674 | 0.775 | 0.777 | 0.713 | 0.286 |
| | **1E-09** | **215** | **0.926** | **0.986** | **0.923** | 8001 | 0.767 | 0.767 | 0.703 | 0.281 |
| | 1E-10 | 219 | 0.917 | 0.977 | 0.915 | 8356 | 0.759 | 0.757 | 0.694 | 0.276 |

[1] Adjusted mutual information (AMI) on homologous superfamily annotation of allergen dataset
[2] Family annotation of allergen dataset
[3] Domain annotation of allergen dataset
[4] Family annotation of ASTRAL SCOPe dataset
[5] Superfamily annotation of ASTRAL SCOPe dataset
[6] Fold annotation of ASTRAL SCOPe dataset
[7] Class annotation of ASTRAL SCOPe dataset
[*] The maximum E-value in MMseqs2 is by default (1E-3)
